# Enhancing Structure-aware Protein Language Models with Efficient Fine-tuning for Various Protein Prediction Tasks

**DOI:** 10.1101/2025.04.23.650337

**Authors:** Yichuan Zhang, Yongfang Qin, Mahdi Pourmirzaei, Qing Shao, Duolin Wang, Dong Xu

## Abstract

Proteins are crucial in a wide range of biological and engineering processes. Large protein language models (PLMs) can significantly advance our understanding and engineering of proteins. However, the effectiveness of PLMs in prediction and design is largely based on the representations derived from protein sequences. Without incorporating the three-dimensional structures of proteins, PLMs would overlook crucial aspects of how proteins interact with other molecules, thereby limiting their predictive accuracy. To address this issue, we present S-PLM, a 3D structure-aware PLM that employs multi-view contrastive learning to align protein sequences with their 3D structures in a unified latent space. Previously, we utilized a contact map-based approach to encode structural information, applying the Swin-Transformer to contact maps derived from AlphaFold-predicted protein structures. This work introduces a new approach that leverages a Geometric Vector Perceptron (GVP) model to process 3D coordinates and obtain structural embeddings. We focus on the application of structure-aware models for protein-related tasks by utilizing efficient fine-tuning methods to achieve optimal performance without significant computational costs. Our results show that S-PLM outperforms sequence-only PLMs across all protein clustering and classification tasks, achieving performance on par with state-of-the-art methods that require both sequence and structure inputs. S-PLM and its tuning tools are available at https://github.com/duolinwang/S-PLM/.

## 1 Introduction

Proteins are fundamental to nearly every biological process and play a crucial role in addressing challenges related to health, energy, and the environment. Consequently, the ability to rapidly and accurately obtain information about target proteins is essential. Computational prediction of protein properties based on their amino acid sequences is a key component of this effort. Protein Language Models (PLMs) have emerged as powerful tools that can uncover underlying patterns within protein sequences and predict various properties by leveraging these patterns [1]. The development and deployment of PLMs currently follow a two-stage paradigm. Initially, PLMs are trained to convert amino acid sequences into mathematical latent representations or embeddings, using techniques such as masked language modeling (MLM) or autoregressive strategies. In MLM, the model predicts masked amino acids based on their surrounding context, while autoregressive models predict subsequent amino acids based on preceding ones [2], [3], [4]. Subsequently, these pre-trained PLMs are adapted using protein property data to perform specific protein-related tasks. Notable PLMs developed under this paradigm include ProtBert [3], ESM2 [5], and ProtGPT [6]. These models have demonstrated promising results in protein property prediction and *de novo* protein design, highlighting their potential for discovering and analyzing new biological insights [7], [8], [9].

Despite these advancements, a significant challenge in developing PLMs is the integration of critical biophysical information into the embeddings. Protein function is intrinsically linked to its three-dimensional (3D) structure and relying solely on amino acid sequences for training PLMs limits their predictive capabilities, especially for tasks that depend heavily on structural information. To address this limitation, some approaches have been developed to enhance sequence-based embeddings with evolutionary or functional data. For instance, methods such as AlphaFold’s Evoformer [10] and MSA Transformer [11] incorporate multiple sequence alignments (MSA) into PLMs. Additionally, ProteinBERT [12] integrates Gene Ontology (GO) [13] annotations into the MLM pre-training framework using a denoising autoencoder trained on corrupted protein sequences and GO annotations, achieving remarkable performance on various protein tasks despite its relatively small model size. These strategies enrich the information contained within sequence-based embeddings, thereby improving the performance of PLMs. However, none of these methods effectively incorporate key structural information into the embeddings.

In response to the need for integrating structural information, emerging approaches aim to develop joint embeddings that combine both sequence and structural data. Chen et al. proposed a protein structure representation method based on self-supervised learning that leverages existing pre-trained PLMs [14]. Similarly, Zhang et al. explored joint protein embeddings by combining ESM2 with three different structure encoders [15], and Hu et al. developed a joint embedding by integrating protein sequences with structural data [16]. These joint embeddings have shown superior performance in many protein property prediction tasks, underscoring the importance of including structural information in protein representations. Nevertheless, a major limitation of these joint embedding models is their requirement for both sequence and structure inputs during inference. Even with tools like AlphaFold, obtaining reliable structures for certain specific proteins remains challenging. Additionally, from a computational standpoint, these methods necessitate additional steps to acquire predicted structures, which increases both time and effort.

To overcome these challenges, we propose an alternative multimodal approach to protein representation learning that does not require simultaneous sequence and structure inputs. Instead of creating joint embeddings, we developed a sequence-based embedding that inherently incorporates structural information. We introduce S-PLM, a 3D structure-aware approach that infuses sequence-based embeddings with structural information by aligning them with structural embeddings derived from 3D protein models through multi-view contrastive learning [17]. A key advantage of S-PLM over existing joint embedding models is that, after training, it only requires amino acid sequences as input during inference, thereby eliminating the overhead associated with using predicted protein structures. This sequence-only input approach enhances computational efficiency and broadens the applicability of the model.

The potential of contrastive learning to enrich sequence-based embeddings is further supported by recent successes. ProtST [18], for example, integrates biomedical text into PLMs by aligning two modalities via a contrastive loss, resulting in enhanced protein property information and superior performance on various protein representation and classification benchmarks. Similarly, the CLEAN model [19] leverages information from EC numbers to improve PLMs’ ability to predict enzyme functions based solely on protein sequences. Building on these successes, S-PLM is designed to accommodate either structure or sequence as input. This study focuses on demonstrating its advantages in encoding protein sequences and enhancing sequence representations for diverse protein prediction tasks using structural information. The sequence encoder of S-PLM is implemented based on the pre-trained ESM2 model, which is scalable and capable of gradually integrating new protein features without forgetting previously learned knowledge [20].

To further enhance the utility of S-PLM, this study builds upon our previous work by introducing a new model, S-PLM2. S-PLM1, introduced in our previous paper [17], uses contact maps and applies protein-level contrastive learning to align sequence and structure embeddings at a global level. In contrast, the newly introduced S-PLM2 incorporates 3D coordinate features and extends to include residue-level contrastive learning, capturing finer structural details. Both models employ efficient fine-tuning techniques to enable targeted modifications for downstream protein prediction tasks without requiring full parameter updates. Efficient fine-tuning strategies such as fine-tuning the top layers, adapter tuning [20], and low-rank adaptation (LoRA) [21] selectively update specific parameters while keeping others fixed. These approaches significantly reduce computational and memory requirements, mitigate issues related to data scarcity, and achieve performance comparable to or better than traditional methods. However, the application of these strategies to PLMs has been limited in existing research [22], [23]. This work explores several efficient fine-tuning strategies for both S-PLM1 and S-PLM2 across various protein prediction tasks. Through these strategies, S-PLM demonstrates competitive performance in gene ontology [13], enzymology number [24], protein secondary structure, protein fold, and enzyme reaction prediction tasks.

## 2 Materials and methods

### 2.1 Structure-aware protein language model (S-PLM)

#### 2.1.1 Overview of the S-PLM framework

An overview of the architectures for both S-PLM1 and S-PLM2 is shown in Figure 1a, showcasing the integration of sequence and structure information through different encoders and the application of contrastive learning at distinct levels. S-PLM1 focuses on aligning sequence and structure representations at the protein level, while S-PLM2 extends this approach to include residue-level contrastive learning. These models aim to capture both the global and fine-grained structural information of proteins, thus enhancing their utility in downstream tasks such as clustering and classification.

**Figure 1.**
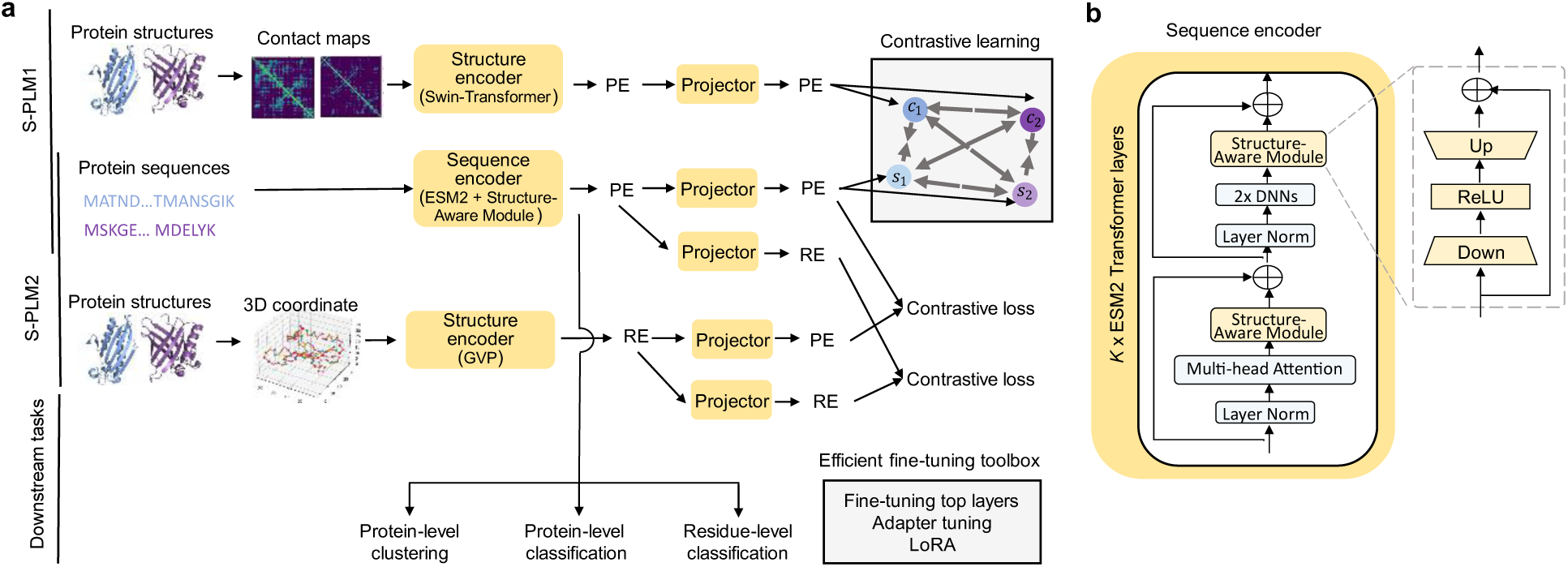
The framework of S-PLM. **a**, The framework of S-PLM1 and S-PLM2. During pretraining, the model inputs the amino acid sequences and contact maps (S-PLM1) or 3D coordinates (S-PLM2) simultaneously. The amino acid sequences of proteins are converted into residue-level embeddings (REs) through a sequence encoder (ESM2 + Structure-Aware Module), while the backbone Cα contact maps (S-PLM1) are converted into protein-level embeddings (PEs) through a structure encoder (Swin-Transformer). For S-PLM2, the 3D coordinates of protein structures are processed through the structure encoder (Geometric Vector Perceptron, GVP) to generate both PEs and REss. With respective projectors, both S-PLM1 and S-PLM2 are trained through contrastive learning on the embeddings from each modality. After pretraining, the sequence encoder that generates the REs before the projector layer is used for downstream tasks, such as protein-level clustering, protein-level classification, and residue-level classification tasks. **b**, Architecture of the sequence encoder. The sequence encoders of S-PLM1 and S-PLM2 are both fine-tuned on the top-K Transformer layers of ESM2, with each layer adding a compact Structure-Aware Module. During pretraining, the ESM2 backbone model is frozen, and only the Structure-Aware Modules are trainable.

In S-PLM1, the model takes as input both the amino acid sequences of proteins and their corresponding structural representations in the form of contact maps. The Swin-Transformer [25] serves as the structure encoder, processing contact maps into protein-level embeddings (PEs), which capture the overall structural configuration of the protein. Simultaneously, the amino acid sequences are processed by the ESM2 model, enhanced with a Structure-Aware Module to incorporate structural information, generating residue-level embeddings (REs). These sequence and structure embeddings are then projected into a unified latent space through independent projectors, utilizing contrastive learning inspired by the SimCLR method [26] and extending the CLIP framework [27]. This ensures that sequence and structure embeddings of the same protein are brought closer together, while maintaining distinct separation between embeddings from different proteins. Moreover, our model accounts for de-alignment within the same modality. The contrastive learning framework in S-PLM^1^ allows for the effective alignment of sequence and structure at the protein level, making it suitable for tasks that require a global understanding of protein structure, such as protein-level clustering and classification.

S-PLM2, on the other hand, introduces a more detailed alignment between sequence and structure by performing contrastive learning at both the protein and residue levels. Here, the structural information is represented by 3D coordinates of the protein, which are encoded through the Geometric Vector Perceptron (GVP) [28]. This structure encoder produces both protein-level and residue-level embeddings (PE and RE, respectively). Like S-PLM1, the sequence embeddings are generated by the ESM2 model with the Structure-Aware Module. The embeddings from both the sequence and structure encoders are then projected into unified latent spaces, where contrastive learning is applied not only at the protein level but also at the residue level. This residue-level contrastive learning enables the model to capture fine-grained details about the spatial relationships between residues, which is crucial for residue-level prediction tasks such as secondary structure prediction. By aligning sequence and structure at multiple levels, S-PLM2 enhances the model’s ability to capture both local and global protein features, making it more effective in tasks that require detailed structural understanding.

The sequence encoder of S-PLM for both versions, depicted in Figure 1b, is built upon the ESM2 Transformer architecture, enhanced with a Structure-Aware Module to integrate structural information into the sequence embeddings. The Structure-Aware Module can be implemented in several ways. One approach involves fine-tuning the pretrained weights of the ESM2 model, allowing the structural information to be incorporated while retaining the original architecture of ESM2. Alternatively, adapter tuning offers a more efficient solution by integrating adapter modules into the top-*K* Transformer layers of ESM2, which are only trained during the pretraining process. These adapter modules, positioned after the multi-head attention projection and following the feed-forward layers, serve as the structure-aware components of the sequence encoder.

Adapter tuning brings several benefits. Firstly, the adapter modules are compact, containing significantly fewer parameters than the full Transformer layers of ESM2, which reduces the training burden. Additionally, this approach enables the model to be continually trained on new protein features (e.g., protein functions) without catastrophic forgetting of previously learned information. This is possible because the ESM2 backbone remains intact and frozen during pretraining, preserving the sequence representation capabilities of the model.

During inference, the S-PLM model is highly flexible, capable of accepting either sequence data or structural data as input. The model produces embeddings suitable for a range of downstream tasks, such as protein-level clustering, protein-level classification, and residue-level classification. In this study, we will focus on generating sequence embeddings from protein sequences by S-PLM, and the pretrained sequence encoder is utilized to produce residue-level embeddings before passing them through projectors for downstream applications. Depending on the task requirements, the entire sequence encoder can either be fully frozen or fine-tuned, ensuring the model’s adaptability. To fully leverage S-PLM’s potential for supervised protein prediction tasks, several efficient fine-tuning strategies, including fine-tuning top layers, adapter tuning, and LoRA, are implemented, all of which are incorporated into the efficient fine-tuning toolbox to optimize performance without excessive computational cost.

#### 2.1.2 Sequence Encoder

The sequence encoder in S-PLM is built upon the ESM2-t33_650M_UR50D model [5], which serves as the backbone for generating contextually rich sequence embeddings. Each protein sequence is tokenized into individual amino acids and processed through 33 Transformer layers, resulting in1280-dimensional embeddings for each amino acid.

To ensure consistency in sequence representation, special tokens are added to the sequences: a BEGIN token (<CLS>) is inserted at the start, an END token ( <EOS>) marks the end, and padding tokens ( <PAD>) are used to standardize sequence lengths for efficient batch processing. The Transformer model generates an embedding for each residue, capturing not only the sequence of residues but also their interdependencies, which are essential for downstream tasks. To optimize computational resources, especially when handling large proteins, sequences exceeding 512 residues are truncated during training. Additionally, S-PLM remains flexible, allowing users to configure the model to process either full-length sequences or truncated ones based on specific task requirements. This flexibility ensures the model can handle a wide range of protein lengths and complexities, from small peptides to larger, more intricate proteins.

The sequence encoder generates residue-level embeddings, which are first projected through two multilayer perceptron (MLP) layers into a 64D representation. The protein-level embedding, on the other hand, is derived by averaging the residue embeddings after excluding the padding tokens, resulting in a global representation of the protein. This global representation is then projected through two additional MLP layers to form a 256D embedding. Both the residue-level and protein-level projections are essential for contrastive learning to ensure that the model can effectively align sequence and structural information. Further details about these projections will be introduced later in the paper.

#### 2.1.3 Structure Encoder: Contact Map with Swin-Transformer

The structure encoder is a crucial component of S-PLM that enables the model to incorporate 3D structural information about proteins. Previously in S-PLM1, we used a contact map-based approach to provide a global view of protein-level residue interactions, utilizing contact maps derived from AlphaFold-predicted structures. However, in this work, we focus on a new approach of S-PLM2 that employs a GVP model, which processes 3D atomic coordinates to capture detailed local structural features at the residue level. This allows for a more precise representation of protein structures. While the contact map method is briefly introduced here for completeness, the GVP model is the primary focus of our current method.

The contact map approach offers a protein-level representation of the structural relationships between residues, simplifying the 3D structure of proteins into a 2D matrix. This makes it well-suited for applying image-processing techniques like Swin-Transformer encoder. To meet the image network’s requirement for three input channels, we transformed the contact map into a three-channel format. Unlike the conventional binary contact map, which assigns a value of 1 when the pairwise distance between Cα atoms is within a defined threshold (indicating residue contact) and 0 otherwise, we converted the contact map into a continuous similarity matrix. In this transformation, *C* represents the elements of the raw contact map. To normalize the raw distance values, any distance in *C* exceeding a threshold *d* (in our case, *d* = Å as used in AlphaFold2) is set to *d*. The transformation formula is:

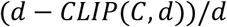

Here, *CLIP*(*C*, *d*) limits values in *C* to a maximum of *d*, ensuring that distances greater than *d* are capped at *d*, while others remain unchanged. The formula scales the distances into a continuous range from 0 to 1, where 1 represents the closest pairwise distance and 0 the farthest. This approach generates a standardized similarity matrix, offering a more consistent representation of the protein’s structural features.

The continuous similarity matrix is then processed by the Swin-Transformer (swinv2-tiny-patch4-window8-256), which extracts hierarchical features at multiple scales. This multi-scale approach enables the model to capture both local and overall architecture of the protein. The resulting protein-level embeddings capture the structural conformation and, like the sequence embeddings, are further refined and reduced to a compact 256-dimensional representation, the same shape as the sequence representation, through two projector layers.

#### 2.1.5 Structure Encoder: GVP with 3D Coordinates

In problems involving protein structure prediction, graph-based models have proven effective due to their ability to represent complex spatial relationships. The GVP model, which leverages a graph structure, excels in representing 3D protein information by incorporating key properties such as rotational invariance and equivariance. In GVP, each residue is represented as a node in a graph, with edges representing spatial relationships. The GVP model can process both scalar and vector features, where scalar features remain invariant, and vector features are equivariant under geometric transformations. These properties ensure that the model maintains geometric consistency during transformations like rotations and reflections, enabling a deeper understanding of how residue arrangements affect the protein’s 3D structure and biological function.

For the node features, both scalar and vector information are utilized. Scalar information is extracted from the geometric properties of the protein structure [28], [29], such as dihedral angles, which are calculated from the directional vectors between three consecutive backbone atoms, with plane normals derived via the cross product. The angle between the normals measures the torsion between residues, and are transformed into sine and cosine representations. Vector node features include both orientation and sidechain information, derived from the backbone’s 3D coordination and the positions of atoms around the backbone. Orientation vectors are calculated by normalizing the differences between consecutive backbone atoms, capturing both forward and backward directions along the protein chain. Sidechain vectors are computed by normalizing the vectors from the Cα atom to the nitrogen and carbon atoms, combining them into a bisector. A perpendicular vector is calculated via the cross product, and a linear combination of both perpendicular and bisector forms the final sidechain vector.

In addition to the node features used in the original GVP model, we incorporate optional Foldseek-derived features to enrich both scalar and vector representations [30]. For scalar features, Foldseek provides cosine similarities between backbone vectors, inter-residue distances, and features encoding the relative sequence distances between residues. For vector features, Foldseek contributes normalized direction vectors and vectors that capture the spatial orientations of residues. These combined Foldseek features further enhance the node representations within the model.

At the edge level, similar to the nodes, both scalar and vector features are incorporated. Scalar features consist of radial basis functions (RBF) to encode pairwise distances between residues [31], and rotary embeddings, which use trigonometric functions to capture relative positions in a rotation-invariant way. Vector edge features represent directional vectors between residues, describing their spatial orientation and distance, essential for maintaining the protein’s structural integrity.

To construct the graph, an optional protein sequence embedding is integrated using two types of embeddings: one-hot encoding, as in the original GVP model, and Atchley factor embeddings [32], which capture key biochemical properties such as polarity, structure factor, molecular volume, amino acid composition, and electrostatic charge. However, an ablation study (Figure 7) showed that incorporating sequence embeddings did not improve model performance.

The network’s architecture processes node and edge features using a GVP-based structure encoder. Node and edge features are passed through the encoder, which captures structural information from the graph. The residue-level embeddings produced by the GVP encoder must pass through a projector before they can be used as input for contrastive learning. These embeddings are transformed through a series of MLP projection layers, reducing their dimensionality to 64, and only after this step is the contrastive loss calculated. This approach is crucial in contrastive learning, as the non-linear projection layers adjust the learned representations to focus on features relevant to the contrastive objective. One explanation is that applying the loss to the projected embeddings helps the model avoid overfitting to specific pre-task details, although some information useful for downstream tasks might be lost in the process. This trade-off allows the model to capture more general features while retaining enough relevant information for effective learning.

Additionally, the entire protein’s representation is generated by pooling the residue embeddings and then passing them through another projector layer to produce a final 256D protein-level embedding, which allows for contrastive learning at the protein level as well.

### 2.2 Multi-view contrastive learning

#### 2.2.1 S-PLM1: protein level contrastive learning

S-PLM1 is a protein-level contrastive learning method that we introduced in a previous work. It aims to align the sequence and structure representations of a protein at the protein level. The method leverages multi-view contrastive learning to bring the sequence and structure embeddings of the same protein closer in the latent space while maintaining clear separation between embeddings from different proteins. By encouraging similar representations (i.e., sequence and structure from the same protein) to cluster together, and dissimilar representations (i.e., from different proteins) to be pushed apart, this approach ensures that the protein’s overall sequence and structure information is well-aligned. Understanding this approach is essential for grasping the residue-level contrastive learning introduced later, as it provides the foundation for aligning sequence and structure representations.

S-PLM1 designed a loss function inspired by the loss used in SimCLR [26]. However, instead of relying on augmented versions of the same input to create different views, it treats the sequence embedding 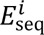 and structure embedding 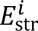 of the same protein as the positive pair. The contrastive loss is defined by the following equation, which measures the similarity between the sequence and structure embeddings of the same protein and contrasts them against embeddings from different proteins in the mini-batch:

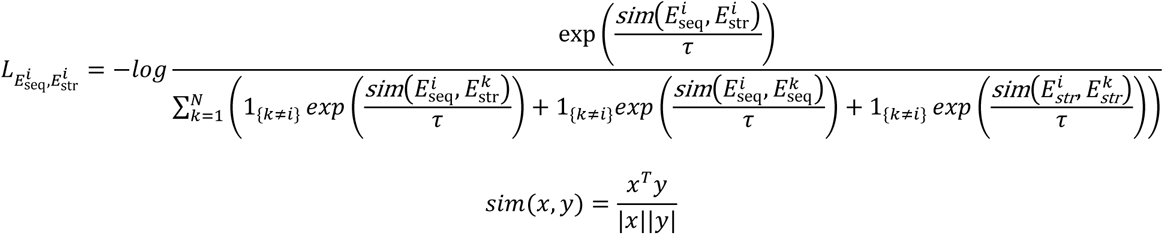

*sim*(*x*, *y*) represents the cosine similarity between embeddings *x* and *y*, while *τ* is the temperature parameter controlling the sharpness of the similarity distribution. The term *k* ≠ *i* ensures that negative examples, corresponding to embeddings from different proteins, are effectively separated in the latent space.

By minimizing this loss, it ensures that the learned representations for a protein’s sequence and structure are well-aligned, capturing both views of the protein in a unified space. At the same time, the model is trained to maintain sufficient distance between the representations of different proteins, preventing confusion across protein types. This multi-view framework leverages both modalities to build a robust representation, highlighting the complementary roles that sequence and structure play in defining protein functions.

#### 2.2.2 S-PLM2: residue level contrastive learning

In S-PLM2, contrastive learning is applied at the residue level to enable more detailed alignment between sequence and structure. As highlighted in a prior study [33], aligning sequence and structure embeddings at the residue level allows for the integration of spatial information from the structure, enhancing the richness of sequence-based representations. This residue-level alignment provides the model with the ability to capture fine-grained structural details, improving its effectiveness in tasks such as secondary structure prediction.

As shown in Figure 2, for each residue, the model computes cosine similarities between the normalized sequence embeddings and structure embeddings, producing a similarity matrix. The similarity matrix is structured so that both its rows and columns follow the same order, with the sequence embeddings listed first, followed by the structure embeddings. Each element in the matrix represents the similarity between the sequence or structure embedding of one residue and the sequence or structure embedding of another residue. However, the diagonal elements of this matrix correspond to the similarity between the same residue’s sequence and structure embeddings, which represents trivial self-similarity. These diagonal elements are not informative for contrastive learning and are therefore discarded.

**Figure 2.**
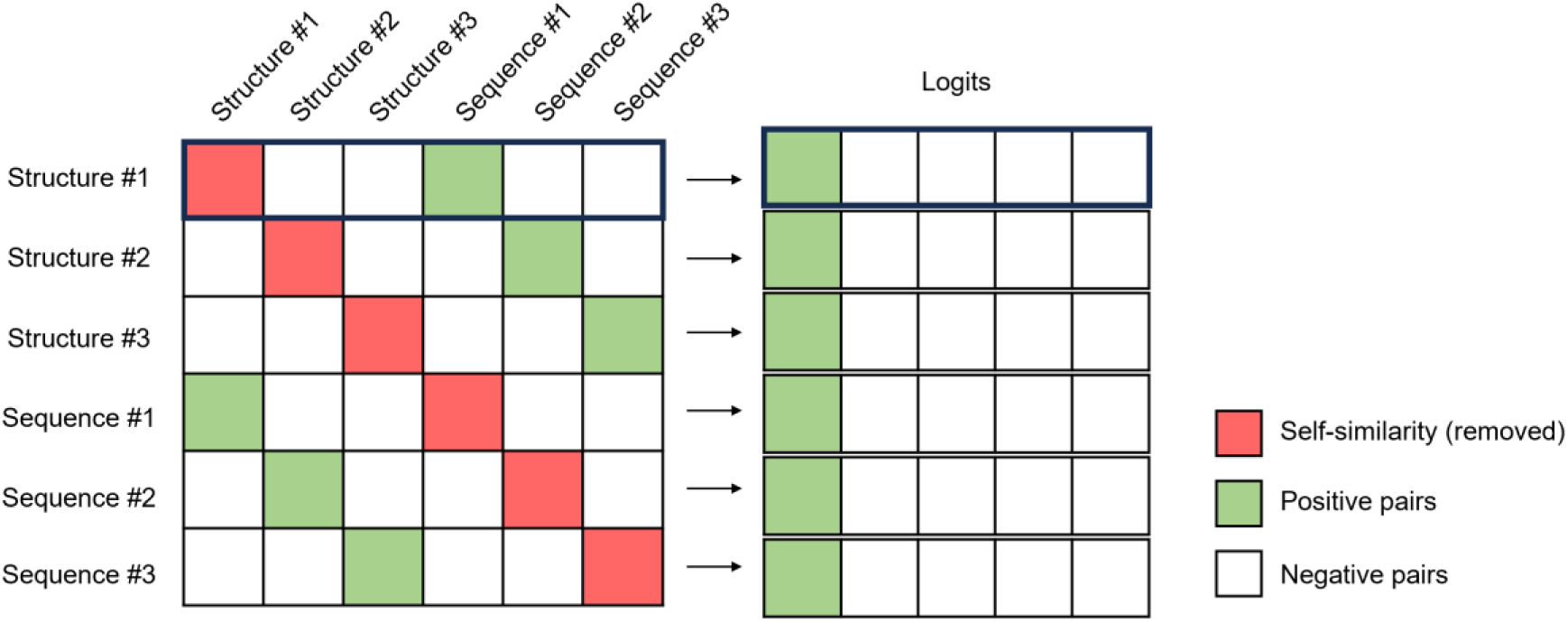
Similarity matrix including self-similarity, positive pairs, and negative pairs in a contrastive learning setting. This diagram only illustrates three residues/proteins as examples for clarity. Rows and columns represent both the structural embeddings (Structure #1, Structure #2, Structure #3) and sequence embeddings (Sequence #1, Sequence #2, Sequence #3). Red cells indicate self-similarity, which is excluded from the contrastive loss calculation. Green cells denote positive pairs, where the structural and sequence embeddings correspond to the same sample and are encouraged to be similar (e.g., Structure #1 with Sequence #1, Structure #2 with Sequence #2, Structure #3 with Sequence #3). White cells represent the remaining negative pairs, which include both pairs from different modalities (e.g., Structure #1 with Sequence #2) and pairs within the same modality (e.g., Structure #1 with Structure #2 or Sequence #1 with Sequence #2).

Once the diagonal elements are removed, the remaining off-diagonal elements represent the meaningful comparisons: positive pairs, which are sequence and structure embeddings from the same residue, while all others are negative pairs. By removing the self-similarities, the model is forced to learn from these more informative comparisons. Positive pairs encourage the alignment of sequence and structure embeddings from the same residue, while negative pairs ensure that embeddings from different residues remain distinct in the latent space.

The selected positive and negative pairs are then used to construct the logits for contrastive learning. These logits are passed into a cross-entropy loss function, where the positive pair logits are expected to have high similarity, while the negative pair logits should have low similarity. The cross-entropy loss applies a softmax function to convert the logits into probabilities, where positive pairs are encouraged to have a probability close to 1, and negative pairs approach 0. This optimization ensures that the sequence and structure embeddings of the same residue are closely aligned, while maintaining separation between embeddings from different residues.

The selected positive and negative pairs are then used to construct the logits for contrastive learning. For each row in the similarity matrix, a single positive pair is combined with multiple negative pairs to form a logit vector. These logits are then passed into a cross-entropy loss function, where the positive pair logits are expected to have high similarity, while the negative pair logits should have low similarity. The cross-entropy loss applies a softmax function to convert the logits into probabilities, where positive pairs are encouraged to have a probability close to 1, and negative pairs approach 0. This optimization ensures that the sequence and structure embeddings of the same residue are closely aligned while maintaining separation between embeddings from different residues.

Formally, the contrastive loss for a positive pair (*i*, *j*) is computed as:

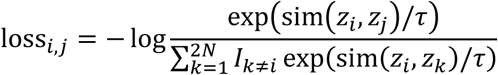

where sim(*z_i_*, *z_j_*) represents the cosine similarity between the embeddings *z_i_* and *z_j_* (the positive pair), *τ* is a temperature parameter that controls the softmax sharpness, and the denominator sums over all pairs, including negative pairs. This formula can be applied from either perspective (i.e., for both (*i*, *j*) and (*i*, *j*) , ensuring that the similarity between structure and sequence embeddings is maximized in both directions. This method of computing the loss is equivalent to the S-PLM^1^.

By minimizing the cross-entropy loss, S-PLM2 effectively captures residue-level relationships, aligning sequence and structure embeddings at a fine-grained level. In addition to residue-level alignment, S-PLM2 also computes global protein-level embeddings by pooling the residue embeddings. This allows for protein-level contrastive learning, complementing the detailed residue-level alignment. Together, the residue and protein-level embeddings provide a robust multi-scale representation, integrating both local structural details and global context. This dual-level alignment enhances the model’s performance in tasks requiring both sequence-structure precision and broader protein-level understanding, such as binding site prediction and protein classification.

### 2.3 Training dataset for contrastive learning and training process

We constructed the training dataset by collecting amino acid sequences from the Swiss-Prot library, and storing them in FASTA format. The corresponding 3D structures of these proteins were sourced from the AlphaFold2 database. In S-PLM1, Cα-Cα contact maps were generated based on the AlphaFold2-predicted structures for each protein. In contrast, S-PLM2 extracted the N, Cα, C, and O atomic coordinates for building a graph representation. From the Swiss-Prot library, we randomly selected 500,000 proteins for training, with an additional 41,500 proteins designated for validation. Given the Swiss-Prot library’s large size, with 542,378 proteins in total, we did not remove similar sequences between the training and validation sets, as this overlap had minimal impact on performance.

During pretraining, proteins longer than 512 residues were truncated to include only the first 512 residues for both sequence and contact map representations. For optimization, we employed two separate SGD optimizers with a momentum of 0.9, one for the sequence backbone and another for the structure backbone. A cyclic learning rate schedule was used, starting at 0.001 and gradually reducing to 0 over 100 steps per cycle. The batch size was fixed at 20, and we applied a weight decay of 0.0005. No gradient accumulation was used. The temperature parameter for contrastive learning was set to 0.05. To stabilize the training process, gradient norm clipping was applied with a threshold of 1.0. Additionally, we adopted mixed precision training [34] to improve computational efficiency and optimize GPU memory usage. All experiments were conducted on a single A100 GPU, running for over 10,000 steps with a consistent batch size of 20. This setup allowed us to effectively balance training complexity with resource constraints, ensuring robust performance throughout the pretraining process.

### 2.4 Efficient fine-tuning on sequence encoder for downstream tasks

Given that the pre-trained models S-PLM1 and S-PLM2 are large in size, full-model fine-tuning is not efficient for downstream tasks due to high computational, memory requirements, and may also lead to suboptimal performance for specific downstream tasks. To address this challenge, we employ efficient fine-tuning strategies to enable targeted modification for downstream protein prediction tasks, applied to the sequence encoder of the S-PLM models, as illustrated in Figure 3. These strategies include fine-tuning top layers, LoRA, and Adapter Tuning.

**Figure 3.**
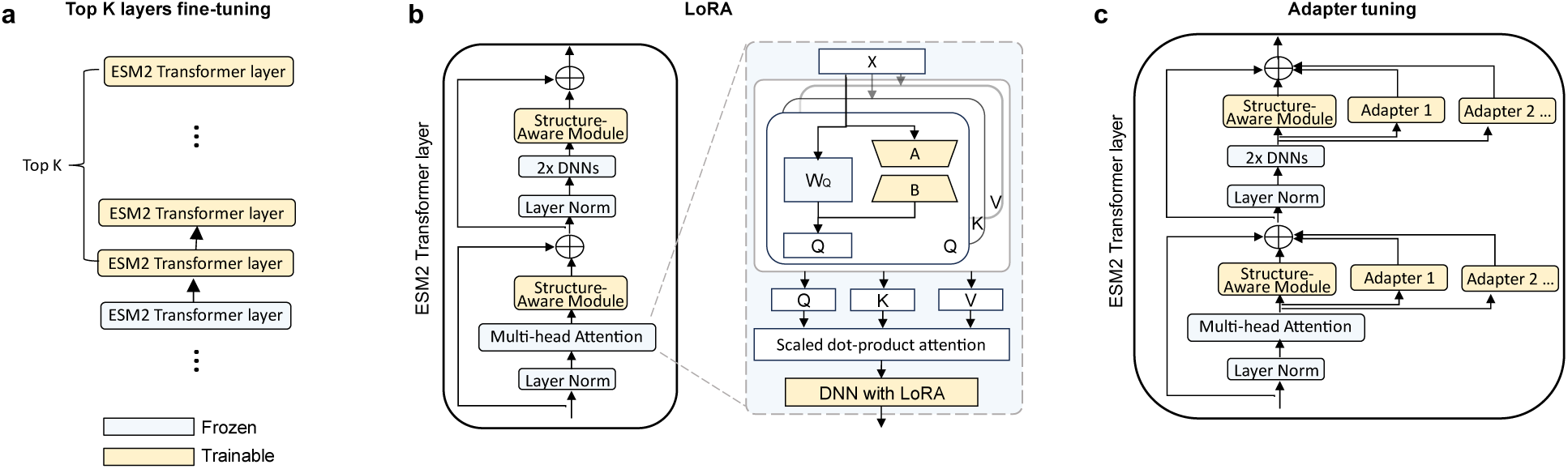
Efficient fine-tuning strategies for the sequence encoder of the S-PLM model. **a**, Fine-tuning top layers. Only the top K Transformer layers are updated, while lower layers remain frozen. **b**, LoRA. Trainable rank decomposition matrices are inserted into the top-K self-attention layers of ESM2. **c**, Adapter Tuning. A list of parallel trainable adapters is added to the Struct-Adapter modules.

The fine-tuning top layers method (Figure 3a) updates only the top *K* Transformer layers of the ESM2 backbone, which consists of 33 layers. By freezing the lower layers, which retain general knowledge from pre-training, and fine-tuning only the top layers, we not only alleviate the computational burden but also reduce the number of trainable parameters. This helps mitigate overfitting to a specific task and prevents the forgetting of pre-trained knowledge. Adjusting the value of *K* provides flexibility, enabling a balance between computational efficiency and task-specific performance.

LoRA (Figure 3b) introduces a more parameter-efficient approach by injecting low-rank matrices into the Transformer’s architecture, while keeping the original pre-trained weight matrices frozen. Instead of updating the entire weight matrix, LoRA represents the update as a low-rank decomposition:

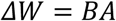

where *B* ∈ *R*^*d×r*^ and *A* ∈ *R*^*r*×*k*^, with *r* ≪ *min*(*d*, *k*). Here, *W*_0_ ∈ *R*^*d*×*k*^ is the original weight matrix. During training, only *A* and *B* are updated, allowing significant reduction in the number of trainable parameters. For an input*x*, the forward pass with LoRA can be expressed as:

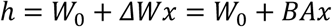

In practice, LoRA scales *ΔW_x_* by 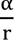 where *α* is a constant controlling the scaling factor. In our implementation, the low-rank decomposition matrices *B* and *A* are applied to the query, key, value, and output projection matrices within the self-attention layers of the top *K* Transformer layers [21] in ESM2. This approach significantly reduces the number of trainable parameters while maintaining the model’s capacity for fine-tuning on new tasks. Here, *K*, *α*, and *r* are hyperparameters in our configuration.

Adapter Tuning (Figure 3c) further optimizes training by inserting adapter modules into the Transformer layers of the ESM2. Unlike the original implementation [20], we only insert adapter modules into top *K* Transformer layers. Each adapter uses a bottleneck structure and a skip connection to compress and reconstruct input features with fewer parameters. Specifically, the adapter compresses the input into a lower-dimensional space and then reconstructs it to match the original dimensions. In our configuration, *K* is a hyperparameter controlling the number of Transformer layers where the adapters are inserted. This method allows scalable fine-tuning for complex bioinformatics tasks while keeping the overall number of trainable parameters low.

For multiple downstream task learning, parallel adapters are employed, where each task-specific adapter is fine-tuned independently while keeping other adapters frozen. This enables efficient fine-tuning across multiple tasks. The overall transformation for each layer is described by:

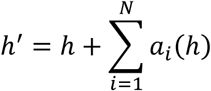

where *h* represents the input features for a Transformer layer, and *N* is the number of parallel adapters. The task-specific transformation of each adapter *a_i_*(*h*) is defined as:

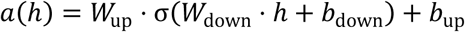

In this equation, *W*_up_ and *W*_down_ are trainable matrices for the up-projection and down-projection, respectively, while *b*_up_ and *b*_up_ are the corresponding biases. The ReLU activation function *σ* is applied to the down-projection.

These efficient fine-tuning strategies, as illustrated in Figure 3, allow us to train large models efficiently while maintaining performance on complex bioinformatics tasks, making the S-PLM model scalable for large-scale protein research.

## 3 Results

### 3.1 Unsupervised protein clustering

3.1.1 Clustering CATH structural hierarchies

CATH [35] is a hierarchical classification system that categorizes protein structures based on their structural and functional features. We assessed the performance of S-PLM by determining its ability to incorporate structural information into the sequence latent space. To emphasize the structural aspects of proteins, we deliberately selected one representative sequence per CATH superfamily and used the CATHS40 dataset, with a maximum sequence identity of 40%. The analysis focused on the class, architecture, and topology levels of the CATH hierarchy, excluding the sequence similarity-driven homologous superfamilies.

We compared the sequence representations generated by S-PLM1 and S-PLM2 with models that rely solely on sequence information, including ESM2 (ESM2_t33_650M_UR50D) — the pretrained PLM on which S-PLM is based — and two other structure-aware models, PromptProtein [36] and ProstT5 [37], which are pretrained by predicting 3D structures or 3D structure tokens from sequences. Figure 4a presents a t-SNE (t-distributed Stochastic Neighbor Embedding) visualization of protein embeddings generated by these models across different CATH hierarchy levels, including Class, Architecture, and Topology. The results indicate that both S-PLM1 and S-PLM2 achieve a clearer and more distinct separation of CATH structural hierarchies compared to the embeddings produced by the other models. This suggests that S-PLM more effectively incorporates structural information into the sequence latent space.

**Figure 4.**
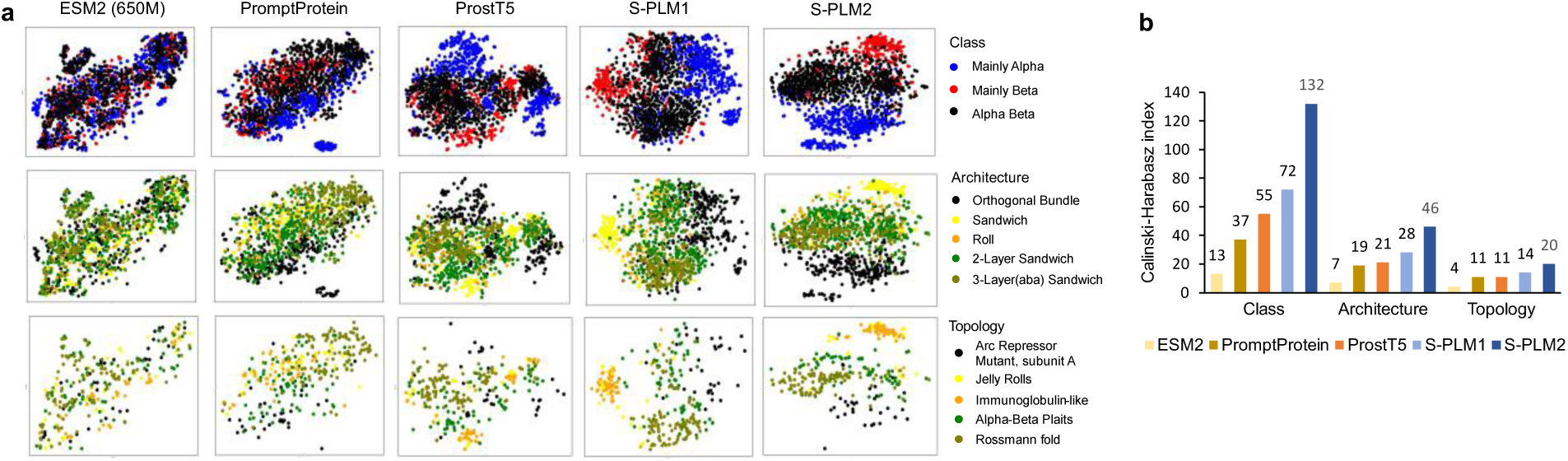
Visualization and benchmark of protein embeddings for CATH structural hierarchies - class, architecture, and topology. **a**, T-SNE visualization of protein embeddings produced by ESM2, PromptProtein, ProstT5, S-PLM1 and S-PLM2. **b**, Utilizing the CHI quantitatively assess the ability of sequence embeddings generated by various models to distinguish CATH structural hierarchies.

The Calinski-Harabasz Index (CHI) [38] measures the ratio between inter-cluster dispersion and intra-cluster dispersion, serving as a metric for cluster validity. To quantitatively assess the ability of sequence embeddings generated by various models to distinguish CATH structural hierarchies, we employed the CHI. In this evaluation, CATH categories were used as ground-truth clusters. As illustrated in Figure 4b, The CHI score of S-PLM1 is approximately 30% higher than that of ProsT5 across the Class, Architecture, and Topology levels, and about 300% higher than that of ESM2. However, S-PLM2 demonstrated even more pronounced gains, with its CHI score being around 140% higher than ProsT5 in the CATH Class and approximately 50% higher in both the CATH Architecture and Topology. Based on the CHI score, S-PLM2 clearly outperforms S-PLM1, suggesting that S-PLM2 is more effective at clustering CATH structural hierarchies, likely due to its directly using of 3D coordinate features and residue-level contrastive learning. Given that these CATH categories are based on protein structures, this analysis strongly supports that the sequence embedding produced by both S-PLM1 and S-PLM2 demonstrate significant awareness of protein structural information, with S-PLM2 offering a more refined representation.

#### 3.1.2 Clustering enzymes

To evaluate the effectiveness of the S-PLM1 and S-PLM2 models in clustering deaminase families, we compared their sequence embeddings with those generated by three other protein language models (PLMs): ESM2, PromptProtein, and ProstT5. The sequence data were sourced from a study by Huang et al., which employed structure-based clustering to identify deaminase functions and novel families undetectable by amino acid sequence analysis alone [39]. We generated 128-dimensional embeddings for each protein using S-PLM1 and S-PLM2, then applied t-SNE to reduce these embeddings to a 2D representation and used K-Means clustering. The adjusted Rand Index (ARI) was calculated by comparing the K-Means clusters with known deaminase family annotations. As shown in Figure 5a and 5b, S-PLM1 achieved an ARI of 0.87, S-PLM2 achieved an ARI of 0.86, outperforming ESM2 (0.63), PromptProtein (0.46), and ProstT5 (0.80).

**Figure 5.**
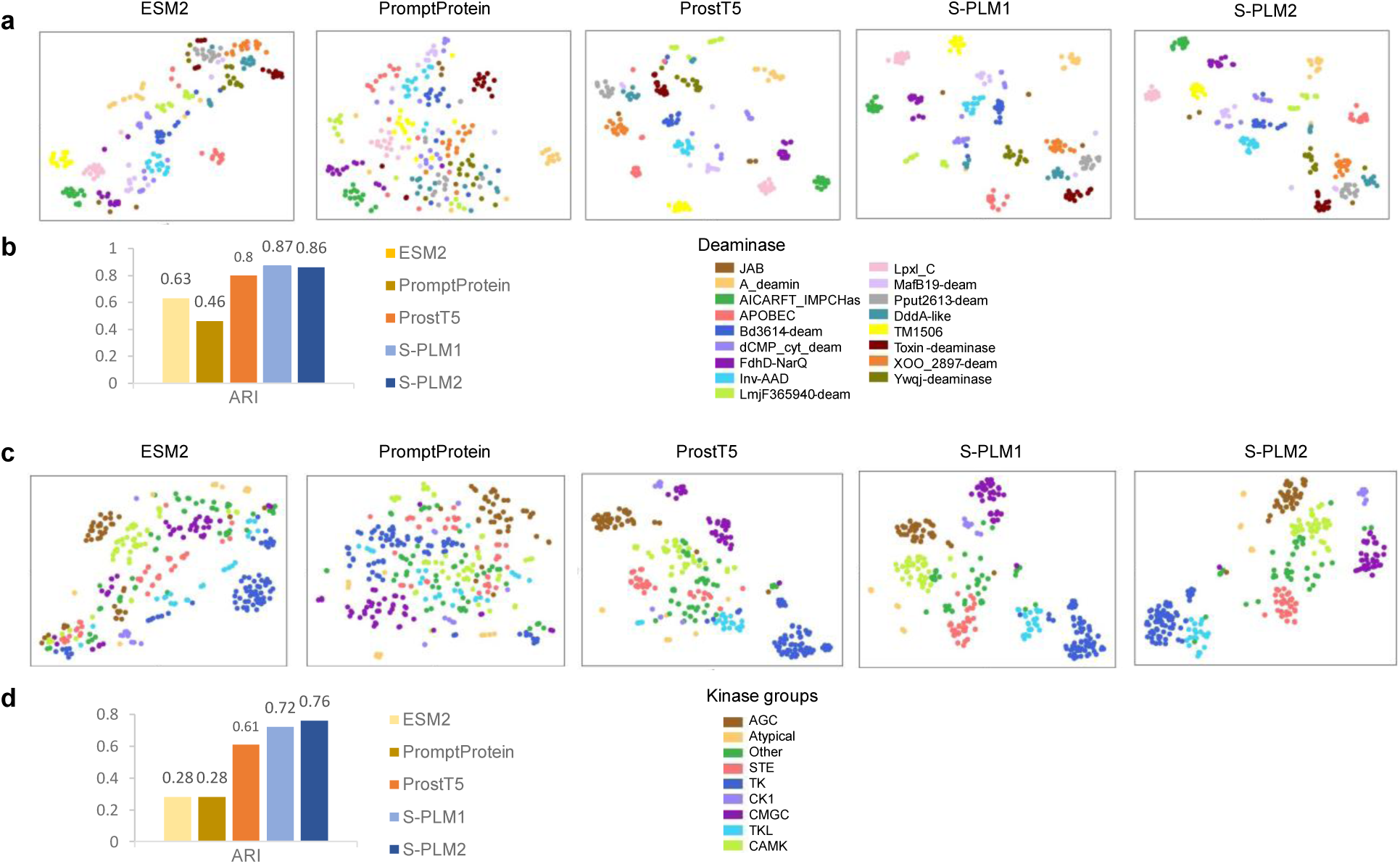
Clustering enzymes. **a**, T-SNE visualization of protein embeddings generated from 242 deaminase proteins using models of ESM2, PromptProtein, ProstT5, S-PLM1 and S-PLM2. **b**, Quantitative benchmarks of the ability of S-PLM’s ability to cluster the deaminase proteins for sequence representation methods. **c**, T-SNE visualization of protein embeddings from human kinase groups using pretrained PLM models of ESM2, PromptProtein, ProstT5, S-PLM1 and S-PLM2. Different kinase groups are distinguished by different colors. **d**, Quantitative benchmarks of the ability of S-PLM1 and S-PLM2 to cluster the human kinases groups compared with other methods mentioned in c. For both types of enzymes, ARI was computed by comparing K-Means clustering assignments with enzyme labels.

We also examined the performance in clustering kinases groups, which facilitate phosphorylation by transferring phosphate groups to proteins. We extracted 336 kinases from GPS5.0, organized into 9 kinase groups, along with their respective kinase domain sequences. Sequence embeddings were generated for each kinase using these domain sequences. For comparison, embeddings were also generated using ESM2, PromptProtein, and ProstT5. As illustrated in the t-SNE plot (Figure 5c) and ARI comparison (Figure 5d), S-PLM1 achieved strong clustering performance for the kinases group, with an ARI of 0.72, while S-PLM2 demonstrated even better performance, with an ARI of 0.78, significantly higher than ESM2 (0.28), PromptProtein (0.28), and ProstT5 (0.61). This superior performance may be attributed to both S-PLM1 and S-PLM2 incorporating structural information essential for characterizing the kinases groups. In summary, S-PLM offers an effective sequence representation for enzyme clustering.

### 3.2 Efficient fine-tuning strategies enhance S-PLM’s performance in selected tasks

To adapt the S-PLM1 and S-PLM2 to various supervised protein prediction tasks, we developed efficient fine-tuning strategies focused on the sequence encoder, including top-layer fine-tuning, adapter tuning, and LoRA. We benchmarked the S-PLM1 and S-PLM2 using various tuning methods across multiple tasks: Gene Ontology (GO) term prediction, Enzyme Commission (EC) number prediction, secondary structure (SS) prediction, protein fold classification, and enzyme reaction (ER) classification, alongside state-of-the-art (SOTA) models. GO term and EC number predictions, as well as protein fold and ER classification, were evaluated as protein-level supervised classification tasks, while SS prediction was assessed as a residue-level task.

For fold classification, we used a dataset of 16,712 proteins classified into 1,195-fold classes from the study [40]. The dataset has three test sets: “Fold,” “Superfamily,” and “Family”. In the “Fold” set, proteins from the same superfamily are excluded from the training data. In the “Superfamily” set, proteins from the same family are excluded from training. In the “Family” set, proteins within the same family are included in the training process. For enzyme reaction classification, a dataset of 37,428 proteins categorized across 384 EC numbers was used, with training, validation, and test splits based on the methodology from the study [41]. GO term prediction and EC number prediction followed the same dataset splits as the study [42], with test sets having up to 95% sequence similarity to the training data. Finally, we adopted Klausen’s dataset as the training set and the CB513 dataset as the test set for secondary structure prediction, maintaining consistency with the reference. The statistics of the dataset for each task are shown in Table 1.

**Table 1:** Dataset statistics of downstream supervised protein prediction tasks.

| Dataset |  | # Train | # Validation | # Test |
| --- | --- | --- | --- | --- |
| Fold classification | Fold | 12,312 | 736 | 718 |
|  | Superfamily | 12,312 | 736 | 1,254 |
|  | Family | 12,312 | 736 | 1,272 |
| Enzyme reaction |  | 29,215 | 2,562 | 5,651 |
| GO term |  | 29,898 | 3,322 | 3,415 |
| EC number |  | 15,550 | 1,729 | 1,919 |
| Secondary structure |  | 8678 | 2170 | 513 |

“#” represents the number of proteins in the subset.

The training, validation, and test sets remained consistent across all tasks, with the only differences being the models and training strategies applied. Detailed results and model descriptions are provided in Table 2.

**Table 2.** Comparing of efficient fine-tuning methods for S-PLM1, S-PLM2, and ESM2 on protein prediction tasks with key model descriptions.

| Method | GO (F<sub>max</sub>) |  |  | #Params | Sequence encoder layers | Classificati<br>on layer |
| --- | --- | --- | --- | --- | --- | --- |
|  | BP | MF | CC |  |  |  |
| S-PLM1-finetune | 0.470 | 0.674 | 0.460 | 42M,40M,<br>40M | finetune top 2 layers,<br>freeze adapter | mean<br>pooling<br>over all<br>residues |
| S-PLM1-finetune | 0.475 | 0.679 | 0.435 |  |  |  |
| ESM2-finetune | 0.473 | 0.677 | 0.450 |  |  |  |
| S-PLM1-adapter | 0.495 | **0.685** | 0.484 | 55M,53M,<br>53M | adapter tuning<br>top 16 layers |  |
| S-PLM2-adapter | **0.498** | 0.684 | **0.485** |  |  |  |
| ESM2-adapter | 0.493 | 0.683 | 0.472 |  |  |  |
| S-PLM1-LoRA | 0.480 | 0.668 | 0.472 | 3M,1M,0.<br>7M | LoRA on top 16 layers,<br>$r = 2, \alpha = 8$ | |
| S-PLM2-LoRA | 0.476 | 0.666 | 0.473 |  |  |  |
| ESM2-LoRA | 0.478 | 0.663 | 0.474 |  |  |  |
| Method | EC (F<sub>max</sub>) |  |  | #Params | Sequence encoder layers | Classificati<br>on layer |
| S-PLM1-finetune | 0.885 |  |  | 40M | finetune top 2 layers,<br>freeze adapter | mean<br>pooling<br>over all<br>residues |
| S-PLM2-finetune | 0.883 |  |  |  |  |  |
| ESM2-finetune | 0.888 |  |  |  |  |  |
| S-PLM1-adapter | 0.875 |  |  | 53M | adapter tuning top 16<br>layers |  |
| S-PLM2-adapter | 0.872 |  |  |  |  |  |
| ESM2-adapter | 0.866 |  |  |  |  |  |
| S-PLM1-LoRA | **0.888** | | | 1M | LoRA on top 33 layers,<br>$r = 2, \alpha = 8$ | |
| S-PLM2-LoRA | 0.883 |  |  |  |  |  |
| ESM2-LoRA | 0.864 |  |  |  |  |  |
| Method | SS (Accuracy%) |  |  | #Params | Sequence encoder layers | Classificati<br>on layer |
| S-PLM1-finetune | 87.48 |  |  | 92M | finetune top 2 layers and<br>all adapter layers | MLP<br>(1280,100,<br>3) on each<br>residue |
| S-PLM2-finetune | **87.49** |  |  |  |  |  |
| ESM2-finetune | 87.21 |  |  |  |  |  |
| S-PLM1-adapter | 87.40 |  |  | 52M | adapter tuning top 16<br>layers |  |
| S-PLM2-adapter | 87.33 |  |  |  |  |  |
| ESM2-adapter | 86.73 |  |  |  |  |  |
| S-PLM1-LoRA | 85.72 | | | 0.8M | LoRA on top 33 layers,<br>$r = 2, \alpha = 8$ | |
| S-PLM2-LoRA | 86.12 |  |  |  |  |  |
| ESM2-LoRA | 87.14 |  |  |  |  |  |
| Method | Protein classification<br>(Accuracy%) |  |  | #Params | Sequence encoder layers | Classificati<br>on layer |
|  | Fold | Superfa<br>mily | Family |  |  |  |
| S-PLM1-finetune | 37.74 | 77.59 | 98.51 | 116M | finetune top 5 layers and<br>all adapter layers | mean<br>pooling<br>over all<br>residues |
| S-PLM2-finetune | **39.42** | 75.44 | 98.58 | 116M |  |  |
| ESM2-finetune | 34.96 | 76.24 | **98.74** | 99M |  |  |
| S-PLM1-adapter | 37.47 | 76.16 | 98.58 | 54M | adapter tuning top 16<br>layers |  |
| S-PLM2-adapter | 38.58 | 76.71 | 98.51 |  |  |  |
| ESM2-adapter | 37.33 | **79.59** | 98.58 |  |  |  |
| S-PLM1-LoRA | 27.04 | 57.42 | 97.34 | 4M | LoRA on top 33 layers,<br>$r = 2, \alpha = 8$ | |
| S-PLM2-LoRA | 27.86 | 64.67 | 98.35 |  |  |  |
| ESM2-LoRA | 27.16 | 67.22 | 97.88 |  |  |  |
| Method | ER (Accuracy%) |  |  | #Params | Sequence encoder layers | Classification layer |
| S-PLM1-finetune | 86.71 |  |  | 23M | Finetune top1 layer and all adapter layers | mean pooling over all residues |
| S-PLM2-finetune | 86.16 |  |  | 23M |  |  |
| ESM2-finetune | 86.53 |  |  | 20M |  |  |
| S-PLM1-adapter | 85.47 |  |  | 53M | adapter tuning top 16 layers |  |
| S-PLM2-adapter | 85.24 |  |  |  |  |  |
| ESM2-adapter | 83.63 |  |  |  |  |  |
| S-PLM1-LoRA | 85.4 | | | 0.8M | LoRA on top 33 layers, $r = 2, \alpha = 8$ | |
| S-PLM2-LoRA | **87.1** |  |  |  |  |  |
| ESM2-LoRA | 84.5 |  |  |  |  |  |

For GO and EC number predictions, seven methods using both sequence and structure inputs— GVP [15], GearNet [43], GearNet-Edge [43], CoupleNet [16], ESM-GearNet [15], PST (finetuned) [44], and SaProt [45]—as well as sequence-only methods ESM2 (650M) [15] and ESM-S [46] were considered for comparison. For the SS task, sequence-only methods, TAPE (Transformer) [47], ProteinBERT [12], ESM-1b [48], and DML [49] were considered for comparison. For protein fold classification, and enzyme reaction (ER) classification, methods using both sequence and structure inputs—GVP, GearNet, GearNet-Edge, CoupleNet were considered for comparison. Their results were taken directly from their respective publications.

Upon comparison as shown in Figure 6, the S-PLM2 demonstrated optimum performance in GO-BP (Fmax: 0.498) and SS task (Accuracy: 0.88), it also exhibits comparable performance in GO-CC (Fmax: 0.494 for CoupleNet, 0.519 for ESM-S, 0.484 for the S-PLM1 and 0.485 for the S-PLM2). At the same time, the S-PLM1 demonstrated optimum performance on GO-MF (Fmax: 0.685), and EC number prediction (Fmax: 0.890 for ESM-GearNet, 0.897 for PST, 0.888 for S-PLM1 and 0.883 for S-PLM2).

**Figure 6.**
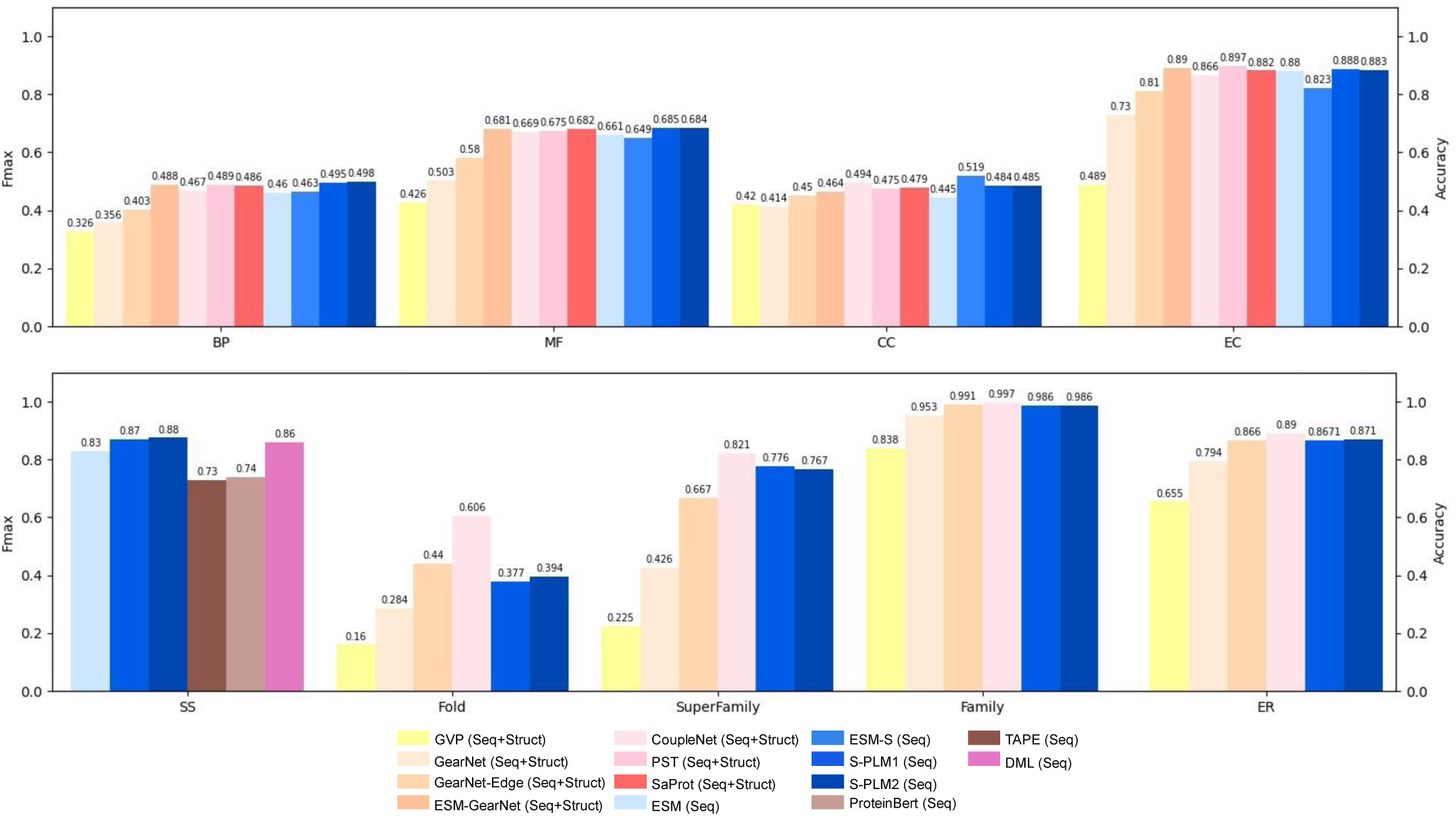
Benchmarking on GO, EC, SS prediction, protein classification, and ER classification. Seq refers to the use of protein sequences as input, while Struct refers to the use of protein structures as input. For GO and EC number predictions, seven methods using both sequence and structure inputs—GVP, GearNet, GearNet-Edge, CoupleNet, ESM-GearNet, PST (finetuned), and SaProt—as well as sequence-only methods ESM2 (650M) and ESM-S were considered for comparison. For the SS task, sequence-only methods, TAPE (Transformer), ProteinBERT, ESM-1b, and DML were considered for comparison. For protein fold classification, and enzyme reaction (ER) classification, methods using both sequence and structure inputs—GVP, GearNet, GearNet-Edge, CoupleNet were considered for comparison.

Among the methods under comparison, GVP and GearNet are designed to capture the invariant and equivariant features of protein structure, whereas GearNet-Edge is a variant of GearNet enhanced with an edge message passing layer. CoupleNet requires the integration of sequence and structure information by deeply coupling them, and ESM-GearNet is a recently proposed variant of GearNet that integrates the sequence representation from PLM with the structure representation through various fusion schemes, where the results with the best fusion scheme were reported in the comparison. SaProt is a newly introduced structure-aware PLM that explicitly utilizes a structure-aware vocabulary to integrate residue tokens with structure tokens derived from protein sequence and structure inputs. In contrast to these methods, our model relies solely on protein sequences. Remarkably, it achieves compatible results to CoupleNet, ESM-GearNet, and PST, or even superior results compared to GVP, GearNet, GearNet-Edge, and SaProt, showcasing the broader practical utility of our model.

Additionally, by employing the same efficient fine-tuning strategies on the base ESM2 encoder with an equal number of trainable parameters, we established a fair comparison among S-PLM1, S-PLM2, and the base ESM2 model. The results are presented in Table 2. In most (17/27) of the one-to-one comparisons, the S-PLM1 encoder outperformed or matched its base ESM2 encoder. In most (16/27) one-to-one comparisons, the S-PLM2 encoder outperformed its base ESM2 encoder. Although we could not surpass ESM2 in all tasks across all training strategies, both S-PLM1 and S-PLM2 models consistently achieved the best overall performance for each task. In summary, our additional Structure-Aware Module on ESM2, along with the new pretraining process, did not diminish ESM2’s original sequence representation capability but rather led to superior performance in general.

### 3.3 Ablation tests

In developing S-PLM, we explored various configurations of sequence encoders and pretraining strategies. This section presents the key findings from our ablation tests for both S-PLM1 and S-PLM2. Specifically, we evaluated the performance of S-PLM1 and S-PLM2 using different trainable layers across various training strategies, including adapter tuning, fine-tuning, and LoRA. The hyperparameter *K* controlled the number of top Transformer layers of the ESM2 base model into which the adapter module was inserted, as well as the number of layers that were fine-tuned or tuned using LoRA. The performance of these configurations was assessed on three protein clustering tasks, including the CATH clustering, deaminase protein clustering, and kinase group clustering. Preliminary training sessions were conducted for 10,000 steps with a consistent batch size of 20. To ensure robustness, each model variant was trained using three different random seeds. The adjusted Rand index (ARI) was computed for each task, with the average ARI score used to provide an overall evaluation.

For adapter tuning, we initially explored larger values of *K* but discontinued further attempts after observing performance declines at higher values. In the case of fine-tuning, the number of trainable parameters increases significantly, which restricted our ability to explore higher values of *K*. As a result, we limited our exploration to values up to 6. For LoRA, larger *K* values were possible, but *K* was capped at 33, as this is the maximum number of layers in the base model.

The results, shown in Figure 7, indicate that increasing *K* for adapter tuning of S-PLM1 generally enhanced clustering performance in the CATH task, particularly in the CATH and topology categories. However, this improvement led to performance degradation in the other two tasks, which was expected. A larger number of adapter layers has a more pronounced effect on the base model, enhancing the structure-aware module’s ability to learn structural features but potentially compromising the sequence representation of the base model. Fine-tuning with *K*=4 led to modest improvements across all tasks, though comparable to the performance gains seen with higher *K* values in adapter tuning. LoRA fine-tuning with larger *K* values demonstrated competitive performance in the CATH clustering task, but similar to adapter tuning, it showed trade-offs in the other clustering tasks, especially when *K* approached the upper limit.

**Figure 7.**
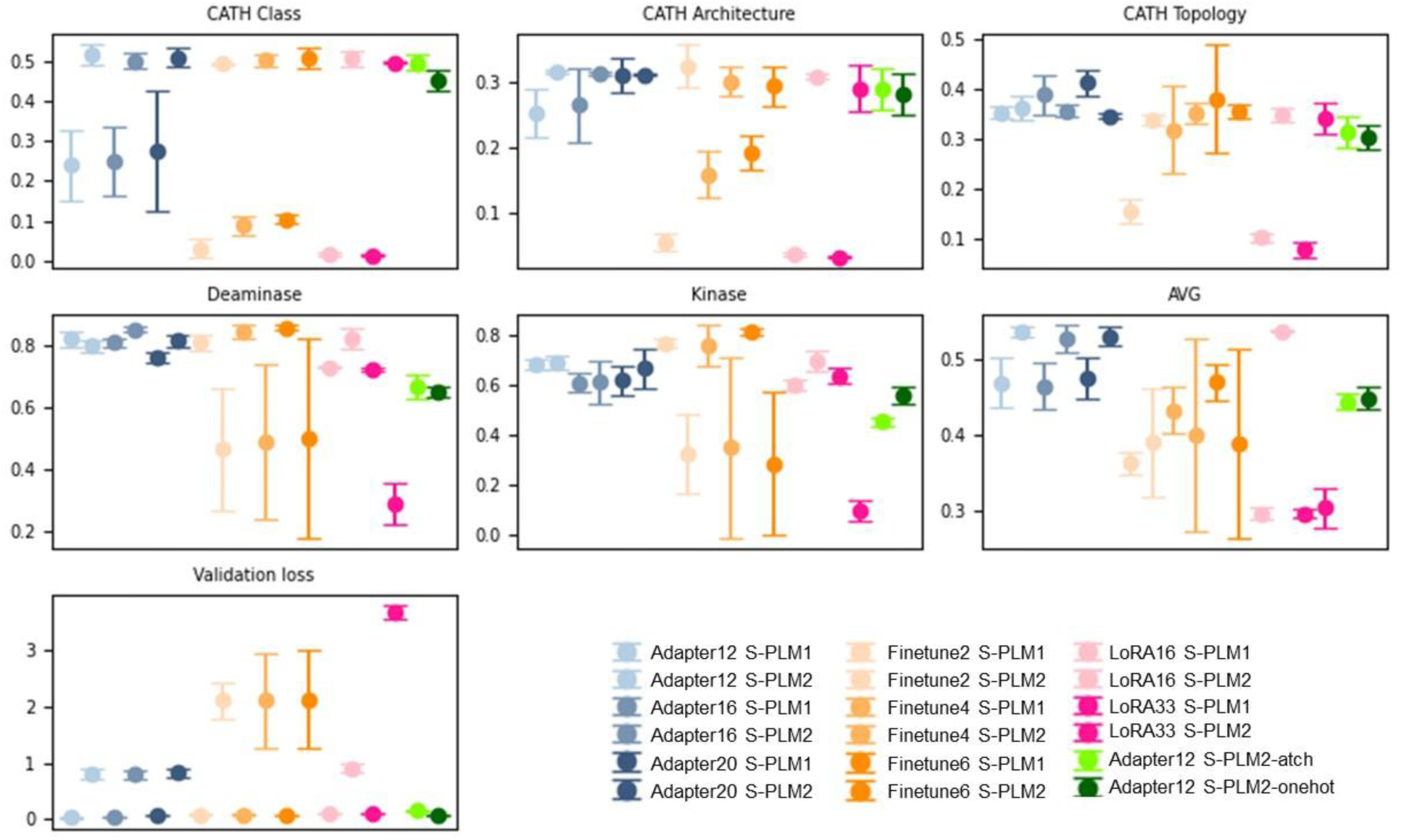
Quantitative evaluation of different configurations of S-PLM1 and S-PLM2. their tuning methods using CATH clustering (Class, Architecture, and Topology), deaminase protein clustering, and kinase group clustering tasks. Performance is measured using Adjusted Rand Index (ARI) by comparing K-Means clustering assignments with true labels. The numbers indicate the specific number of ESM2 Transformer layers where the adapter module is inserted, or the number of layers fine-tuned or tuned using LoRA for each method.

Based on these findings, we ultimately selected adapter tuning with *K*=16 for S-PLM1 and *K*=12 for S-PLM2, achieving a balance between structural feature learning and the preservation of sequence representation performance. For S-PLM2, we introduced additional parameters to manage the usage of sequence embedding in the GVP model, specifically exploring the Atchley factor embedding and one-hot encoding. However, as demonstrated in Figure 7, including these sequence embeddings did not improve performance and, in fact led to worse outcomes compared to not using sequence embeddings.

## 4 Discussion

In this work, we introduced two versions of S-PLM (S-PLM1 and S-PLM2), structure-aware protein language models pretrained using contrastive learning between protein sequences and their 3D structures. S-PLM1 uses contact maps and applies protein-level contrastive learning to align sequence and structure embeddings at a global level, while S-PLM2 uses 3D coordinate features and extends to include residue-level contrastive learning, capturing finer structural details. Both models employ efficient fine-tuning techniques to enable targeted modifications for downstream protein prediction tasks without requiring full parameter updates. Unlike joint-embedding methods, S-PLM independently encodes sequences and 3D structures, enabling structure-aware predictions using only sequence data.

The results of our study demonstrate S-PLM’s ability to incorporate structural information into sequence representations, excelling in both protein clustering and supervised protein prediction tasks. In unsupervised clustering of CATH structural hierarchies and enzymes, both S-PLM1 and S-PLM2 showed clearer separations of protein structures compared to sequence-only models like ESM2 and structure-aware models like PromptProtein and ProstT5. Notably, S-PLM2 outperformed S-PLM1 in CATH clustering, which relies heavily on structural features. In enzyme clustering tasks, both models achieved comparable results and significantly outperformed other models, while S-PLM2 surpassed S-PLM1 in kinase clustering. These findings highlight GVP’s strength in encoding intricate 3D structure features, underscoring its effectiveness as a protein structure encoder.

In supervised protein prediction tasks, S-PLM achieved competitive results without using 3D structural inputs, performing similarly to state-of-the-art models that require both sequence and structure. With efficient fine-tuning strategies like fine-tuning, adapter tuning, and LoRA, S-PLM models excelled in tasks such as GO term prediction, EC number prediction, secondary structure prediction, protein classification, and enzyme reaction classification.

We also observed that different tuning strategies performed optimally for different tasks. The variation in performance is likely due to the number of trainable parameters and the level of model adaptation provided by each strategy. Fine-tuning the top layers generally exposes more parameters, while LoRA, despite having fewer trainable parameters, impacts multiple layers, allowing it to capture complex patterns better suited for certain tasks. The nature of the task also plays a role—some tasks may benefit from fine-grained adaptation of lower layers, while others require substantial changes to higher-level representations.

In conclusion, by incorporating a Struct-Aware Module into the pretrained ESM2 model, S-PLM retains ESM2’s sequence representation capabilities while effectively integrating structural information. S-PLM and its efficient fine-tuning strategies offer a powerful and flexible solution for protein analysis and prediction tasks.

## Acknowledgments

We thank Dr. Jin Chen for providing input on the development of the structure-aware protein language models. D.W. and D.X. are partially supported by the National Institutes of Health (grant R35GM126985). Q.S. and D.X. would like to acknowledge the National Institutes of Health (grant R01LM014510). The computation for this work was partially performed on the high-performance computing infrastructure provided by Research Computing Support Services at the University of Missouri. We also thank the University of Kentucky Center for Computational Sciences and Information Technology Services Research Computing for their support and use of the Morgan Compute Cluster and associated research computing resources. This work also used Delta-GPU at NCSA through allocation CIS230053 from the Advanced Cyberinfrastructure Coordination Ecosystem: Services & Support (ACCESS) program, which is supported by National Science Foundation grants #2138259, #2138286, #2138307, #2137603, and #2138296.

